# Arsenite exposure inhibits histone acetyltransferase p300 for attenuating H3K27ac at enhancers in low-dose exposed mouse embryonic fibroblast cells

**DOI:** 10.1101/278424

**Authors:** Yan Zhu, Yanqiang Li, Dan Lou, Yang Gao, Jing Yu, Dehui Kong, Qiang Zhang, Yankai Jia, Haimou Zhang, Zhibin Wang

## Abstract

Epidemiological investigations have validated the positive relationships between arsenic in drinking water and several cancers, including skin, liver and lung cancers. Besides genotoxicity, arsenic exposure-related pathogenesis of disease is widely considered through epigenetic mechanisms; however, the underlying mechanistic insight remains elusive. Herein we explore the initial epigenetic changes via acute low-dose arsenite exposures of mouse embryonic fibroblast (MEF) cells and Dot1L knockout MEF (Dot1L-/- for abbreviation) cells. Our RNA-seq and Western blot data demonstrated that, in both cell lines, acute low-dose arsenite exposure abolished histone acetyltransferase p300 at the RNA level and subsequent protein level. Consequently, p300-specific main target histone H3K27ac, a marker separating active from poised enhancers, decreased dramatically as validated by both Western blot and ChIP-seq analyses. Concomitantly, H3K4me1 as another well-known marker for enhancers also showed significant decreases, suggesting an underappreciated crosstalk between H3K4me1 and H3K27ac involved in arsenite exposure. Significantly, arsenite exposure-reduced H3K27ac and H3K4me1 inhibit the expression of genes including *EP300* itself and *Kruppel Like Factor 4(Klf4)*, a tumor suppressor gene. Collectively, our investigations identify p300 as an internal bridging factor within cells to sense external environmental arsenite exposure to alter chromatin, thereby changing gene transcription for disease pathogenesis.

## 1. Introduction

Arsenic, as a ubiquitous element, ranks the 20th most abundant element in the earth’s crust (Mandal and Suzuki, 2002). Human exposure to environmental arsenic occurs primarily via ingestion of inorganic arsenic contaminated water. It is estimated that over 200 million individuals worldwide are exposed to drinking water containing arsenic above 10 μg/L, the safety standard set by the World Health Organization (Argos *et al.*, 2014). Through numerous epidemiological investigations, accumulating evidence demonstrate the positive relationship between inorganic arsenic and carcinomas, including both skin and internal cancers. In Wisconsin, USA, residents who were over 35 years old and consumed arsenic-contaminated water for more than a decade suffered a higher prevalence of skin cancer than control group (Knobeloch *et al.*, 2006). A significant dose-response relationship between arsenic concentrations and other tumors was found in Taiwan, including bladder, kidney, and lung cancer in both sexes, and tumors of prostate and liver in males (Wu *et al.*, 1989). However, the exact pathogenic mechanisms underlying inorganic arsenic exposure remain to be determined. Inorganic arsenic shows genotoxicity through several aspects, including chromosomal aberration (Oya-Ohta *et al.*, 1996), genomic instability (Bhattacharjee *et al.*, 2013) and p53 dysfunction (Hamadeh *et al.*, 1999). Besides, the role of inorganic arsenic in epigenome has also been initially explored, including DNA methylation, microRNA, and histone modification. The methylation of CpG dinucleotides at the 5’ position on the pyrimidine ring to form 5-methylcytosine (5-mC), which can disrupt transcription by blocking the binding of transcription factors and attracting methyl-binding proteins that initiate chromatin compaction and bring about gene silencing (Lunnon and Mill, 2013). So methylation results in gene silencing, while de-methylation causes gene activation (Tellez-Plaza *et al.*, 2014). S-adenosyl-methionine (SAM) is essential for methylation of inorganic arsenic to detoxication, and it is also the methyl-donor required by DNA methyltransferases. And it was validated that arsenic exposure was associated with hypo-methylation of DNA and facilitates oncogenes expression (Zhao *et al.*, 1997). Besides, hypermethylation of *p53* and *p16* promoters were found in people exposed to arsenic. Histone modification (including histone acetylation and methylation), a covalent post-translational modification, plays an integral role in gene transcription (Allfrey and Mirsky, 1964; Dai H, 2014). For instance, chronic arsenite exposure decreased H4K16ac in UROtsa cells (Jo *et al.*, 2009). However, systematic characterizations to identify effector histone acetyltransferases and altered genomic loci with reduced histone acetylations remain to be done.

For histone acetylation, it is regulated by two group of enzymes, histone acetyltransferases (HATs) and deacetylases (HDACs), with antagonizing functions (Wang *et al.*, 2009). Among members of HATs, p300 and CBP are homologous. They are also global transcriptional co-activators (Ogryzko *et al.*, 1996), and both of them play integral roles in regulation of gene transcription via at least two (not necessarily mutual exclusive) mechanisms. In scenario 1, p300 acts as associated partners of transcription factors (TFs). p300 is recruited by these TFs including p53 to acetylate nucleosomes at promoters and/or enhancers for opening chromatin, thereby facilitating the recruitment of RNA polymerase II and other general transcription factors (Guermah *et al.*, 2006; Wang *et al.*, 2009). For example, our data demonstrate that the increases of histone H3K9ac and H4K16ac at transcription start sites, via inhibition of HDACs with HDAC inhibitor drugs, facilitate the RNA polymerase II recruitment. In scenario 2, p300 functions through acetylation of TFs themselves, thereby impacting the function of TFs. For example, p300 directly acetylates exposure-relevant TF Nrf2. Acetylations of multiple lysine residues of Nrf2 increase the binding of Nrf2 to promoters (Sun *et al.*, 2009). This increased Nrf2 binding presumably contributes to the regulation of Nrf2-regulated gene pathways. In addition, the loss of p300 functions or simple reduction of p300 protein levels has linked to human diseases. For example, DNA mutations of p300 are linked to a small percentage of patients of Rubinstein-Taybi syndrome (Roelfsema *et al.*, 2005). Such mutations lead to either truncated non-functional p300 molecules, or the loss of one copy of *EP300* gene in each cell, thereby reducing the p300 amount by half. While the reduction of p300 amount is linked to development of Rubinstein-Taybi syndrome, the exact underlying mechanism is not clear. Lastly, it seems that p300/CBP specifically mediate H3K18ac and H3K27ac, whereas two other homologous HATs, GCN5 and PCAF, mediate H3K9ac (Jin *et al.*, 2011).

Herein our analyses reveal that sodium arsenite exposure-related health problems may be linked to the reduction of the amount of p300 in exposed cells. We found that acute arsenite exposure in low doses reduced the expression of p300 at both the mRNA levels via RNA-seq and qRT-PCR analyses and the protein levels via Western blotting. The consequence of reduced p300 (i.e., reduction of H3K27ac, an active marker for enhancers) was further identified. In addition, we examined the expression of tumor suppressor genes. Collectively, our data reveal an intriguing epigenetic mechanism that arsenic exposure may impact human health through altering functions of p300 and following histone acetylation modifications it mediated.

## 2. Materials and methods

### 2.1. Cell culture and arsenite exposure

MEF and Dot1L-/-are diploid and nontumorigenic. Both cells were cultured in high glucose Dulbecco’s Modified Eagle Medium (DMEM) with 10% (v/v) fetal bovine serum (FBS) in an incubator at 37 °C with 5% CO_2_. At approximately 80% confluent, cells were treated with fresh medium containing 0, 2, 5, 10 μM sodium arsenite for 24 h. After exposure, cells were collected for following experiments.

### 2.2. Western Blot

Total protein extracts were prepared from cells and loaded into each lane of a 4-12% SDS-PAGE gel as described before (Wang *et al.*, 2008). After electrophoresis, total proteins were transferred onto a nitrocellulose membrane. Membranes were blocked in TBS-T buffer (20 mM Tris base, 150 mM NaCl, 0.05% Tween 20, pH 7.4) containing 5% nonfat milk at room temperature for 2-4 h prior to the addition of primary antibody and incubation at 4°C overnight. Nrf2 antibody (Santa Cruz, sc-13032, Lot # C1416), p300 antibody (Santa Cruz, sc-584, Lot # C0805), H3K4me1 antibody (Abcam, ab8895, Lot # GR243233-1) and H3K27ac antibody (Abcam, ab4729, Lot # GR103379-1) were used to detect each protein. TBS-T-washed membranes were incubated with goat anti-rabbit (Santa Cruz, sc-2004, Lot # A1416) or goat anti-mouse (Santa Cruz, sc-2005, Lot #J2215) secondary antibodies in TBS-T–5% nonfat milk and washed with TBS-T, and membrane-bound antibody was visualized in a chemiluminescence assay. Beta-actin (Sigma, A5441, Lot # 030M4788) was used for normalization.

### 2.3. mRNA extraction and characterization

Total RNAs from one exposed and three control groups of two cell lines were isolated and quality controlled with protocols from Illumina. A total amount of 2 μg RNA per sample was used as input material for library construction. Ribosomal RNA (rRNA) was removed through Epicentre Ribo-zero rRNA Removal Kit (Epicentre, USA) according to the manufacturer’s instructions. Strand-specific sequencing libraries were generated through the dUTP method by using the resulting RNA from NEBNext Ultra Directional RNA Library Prep Kit for Illumina (NEB, USA) following manufacturer’s recommendations. RNA-seq was performed on an Illumina Hiseq 2000 platform and 100 bp paired-end reads were generated according to Illumina’s protocol.

### 2.4. Whole-genome gene expression analysis

The adapter sequences were removed from the raw sequencing data and the individual libraries were converted to the FASTQ format. Sequence reads were aligned to the mouse genome (mm10) with TopHat2 (v2.0.9) and the resulting alignment files were reconstructed with Cufflinks (v2.1.1) and Scripture (beta2). For mRNA analyses, the RefSeq database (Build 37.3) was chosen as the annotation references. The read counts of each transcript were normalized to the length of the individual transcript and to the total mapped fragment counts in each sample and expressed as fragments per kilobase of exon per million fragments mapped (FPKM) of mRNAs in each sample. The mRNA differential expression analyses were applied for exposed and control groups. An adjusted P value <0.05 (Student’s t-test with Benjamini-Hochberg false discovery rate (FDR) adjustment) was used as the cut-off for significantly differentially expressed genes. Differentially expressed genes (DEGs) were analyzed by enrichment analyses to detect over-represented functional terms present in the genomic background. Gene ontology (GO) analysis was performed using the GO-seq R package, in which gene length bias was corrected.

### 2.5. Gene expression validation by quantitative Reverse Transcription-PCR (qRT-PCR)

Total RNAs were isolated and quality controlled using protocols from Illumina, Inc. All cDNAs were synthesized using Superscript III (Invitrogen; Thermo Fisher Scientific, Inc., Waltham, MA, USA). Five over□ and under-expressed genes were selected for qRT□PCR assay. All qRT□PCR primers are presented in Table 1. All qPCR reactions were performed on a Roche Lightcycler 480 PCR system (Roche Applied Science, Penzberg, Germany) using Toyobo Thunderbird SYBR RT□qPCR Mix (Toyobo Life Science, Osaka, Japan), with three technical repeats. The amplification procedure was as follows: 95 °C for 5 min, followed by 40 cycles of 95 °C for 10 sec and 60 °C for 20 sec. Relative quantification of target genes was performed using the 2-ΔΔCT method with GAPDH as a reference gene (Livak and Schmittgen, 2001). Pearson correlation was used to calculate the association between RNA sequence and qRT□PCR (Hane *et al.*, 1993).

**Table 1.**
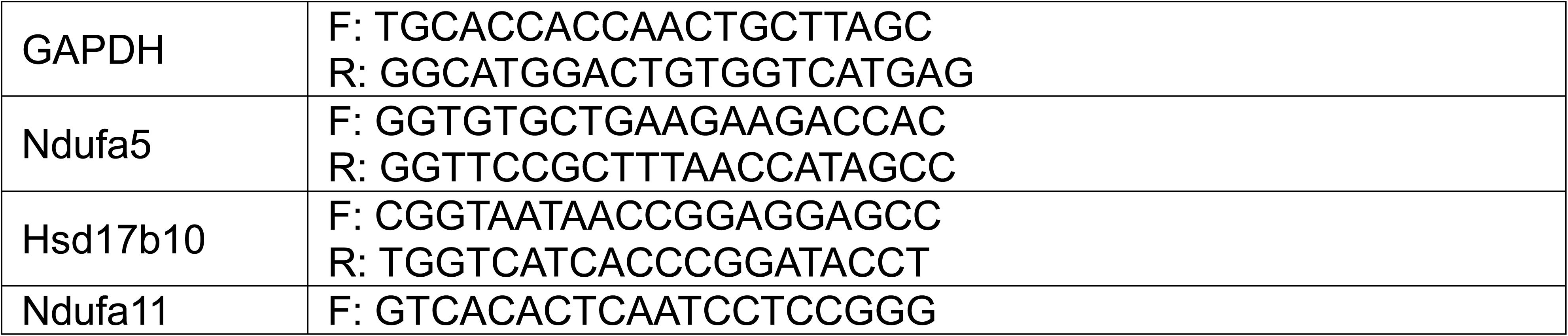
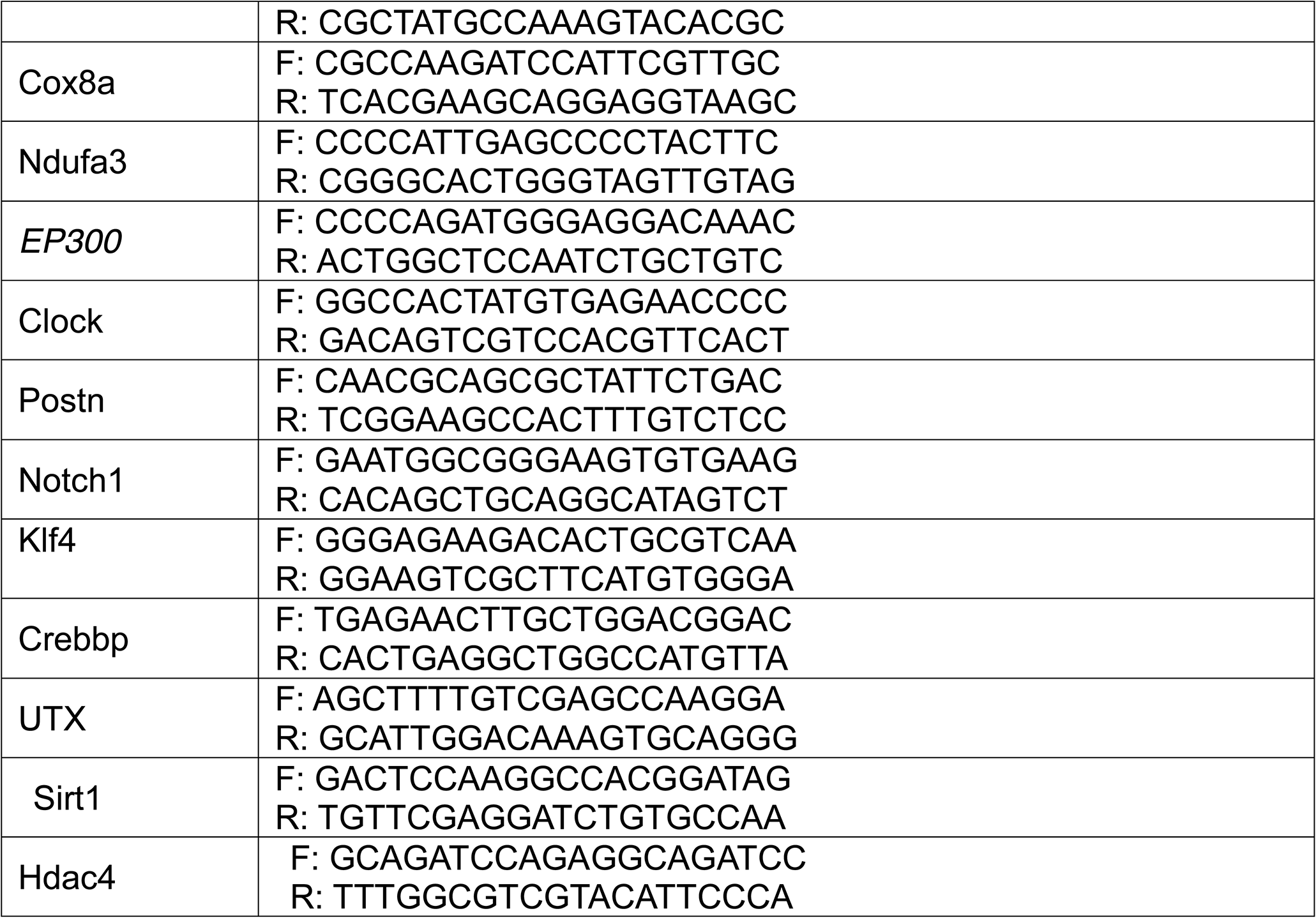
Primers used for qRT-PCR

### 2.6. Chromatin immunoprecipitation sequencing (ChIP-Seq)

Chromatin immunoprecipitation (ChIP) was performed as described previously (Wang *et al.*, 2009). In brief, cells were crosslinked with 1% formaldehyde for 10 min at room temperature and the reaction was quenched with 125 mM glycine. After chromatin fragmented to 200 to 500 bp by sonication, chromatin templates from 20 million cells were used for ChIP experiment. Samples were immunoprecipitated with 2-4 μg of H3K4me1 antibody or H3K27ac antibody overnight at 4 °C followed by washing steps. After reverse crosslinking, the ChIP DNA fragment were purified and repaired using End-It^TM^ DNA End-Repair Kit (Epicentre) followed by treatment with Taq polymerase to generate single-base 3’ overhangs used for adaptor ligation. Following ligation of a pair of Illumina adaptors to the repaired ends, the ChIP DNA was amplified using the adaptor primers. Amplified DNA fragments around 300 bp were isolated from agarose gel prior to sequencing.

### 2.7. Identification of binding regions and peaks

For ChIP-seq of H3K4me1 and H3K27ac, reads were mapped to the mm10 genome using Bowtie aligner allowing up to two mismatches (Langmead, 2010). Only the uniquely mapping reads were used for further analysis. Macs2 call peak program was used to call the peaks and the macs2 bdgdiff was used to identify the difference between MEF and Dot1L (Zhang *et al.*, 2008).

### 2.8. ChIP–qPCR validation

Five genes’ promoters were selected for qRT-PCR assay and presented in Table 2. All qPCR reactions were performed on a Roche Lightcycler 480 PCR system (Roche Applied Science, Penzberg, Germany) using Toyobo Thunderbird SYBR RT□qPCR Mix (Toyobo Life Science, Osaka, Japan), with three technical repeats. The amplification procedure was as follows: 95 °C for 5 min, followed by 40 cycles of 95 °C for 10 sec and 60 °C for 20 sec. The cycle threshold (Ct) of samples was used for calculation. Each value was normalized to the percentage of Input DNA by using IP/INPUT= 2 ^CtINPUTDNA-CtIPDNA^ (Alamdar *et al.*, 2017).

**Table 2.**
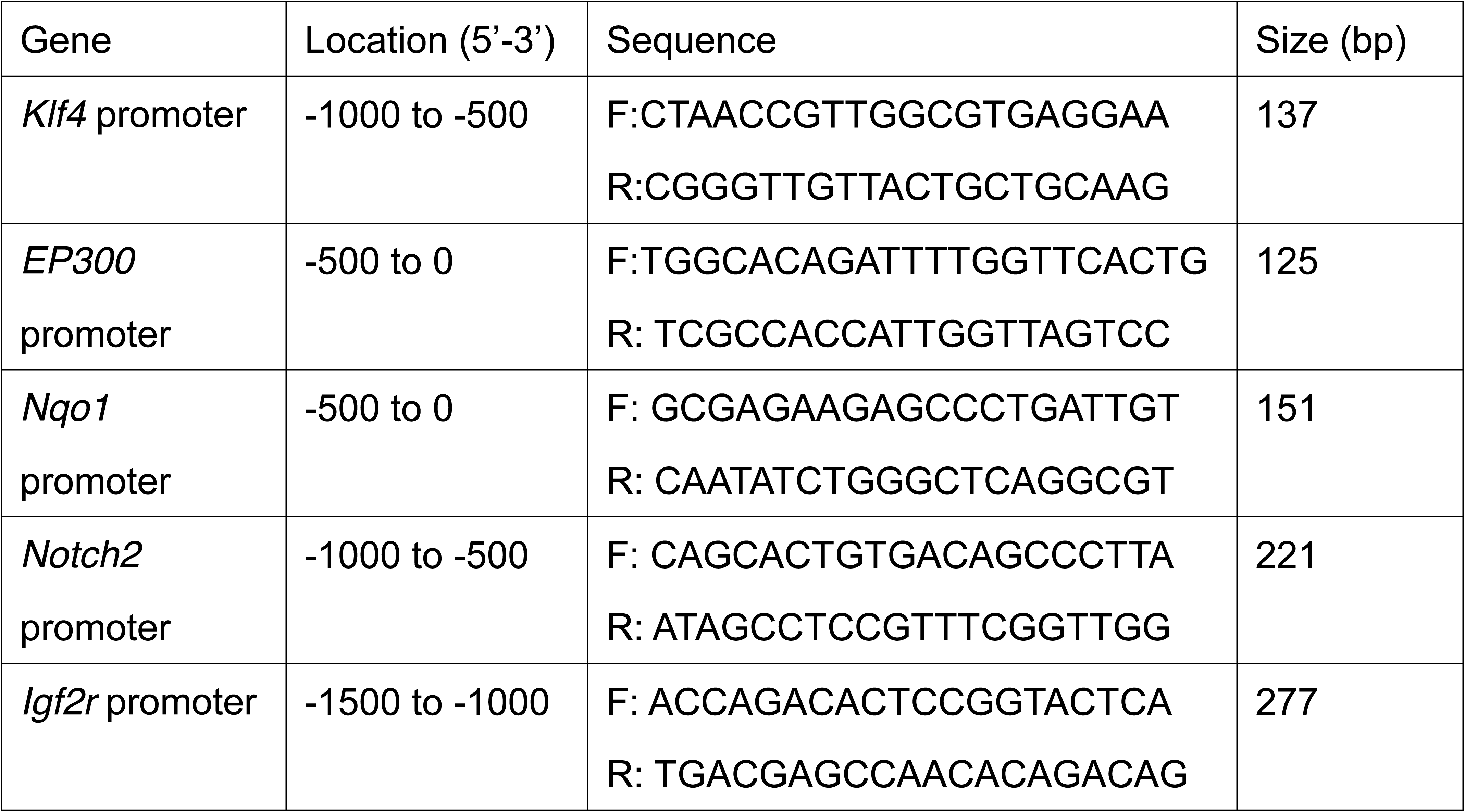
Primers used for ChIP–qPCR validation

### 2.9. Statistical analysis

Results were presented in Means ± standard error of the mean (SEM). Student’s t-test was used to examine difference between two groups and ANOVA was used for multi-groups. * means p<0.05, **means p <0.01.

## 3. Results

### 3.1. Identification of a low dose exposure condition

Toward a low dose exposure condition for epigenetic insights of arsenite exposure, we set up exposure experiments and used Nrf2 expression as a readout to characterize. Based on previous reports, we started our exposure experiments with two time points (6 and 24 h) and four concentrations (0, 2, 5, and 10 μM). Without arsenite exposure, there was limited amount of Nrf2 expressed in both MEF and Dot1L-/-cells (Fig. 1A and 1B). After 6 h treatment, the amount of Nrf2 showed dose-effect relationships with the concentrations of sodium arsenite in both cell lines (Fig.1C). In addition, there was a significant difference between 2 μM and 5 μM groups. Results after 24 h exposure (Fig. 1D and 1E) were similar to 6 h treatment. However, compared with 6 h groups, which showed same results in both cells, 24 h groups (Fig. 1F) showed different pattern of Nrf2 concentrations, which might be caused by *Dot1L* lacking. Collectively, we consider 5 μM and 24 h appropriate for our mechanistic exploration and choose this exposure condition for further experimental characterizations.

**Fig. 1.**
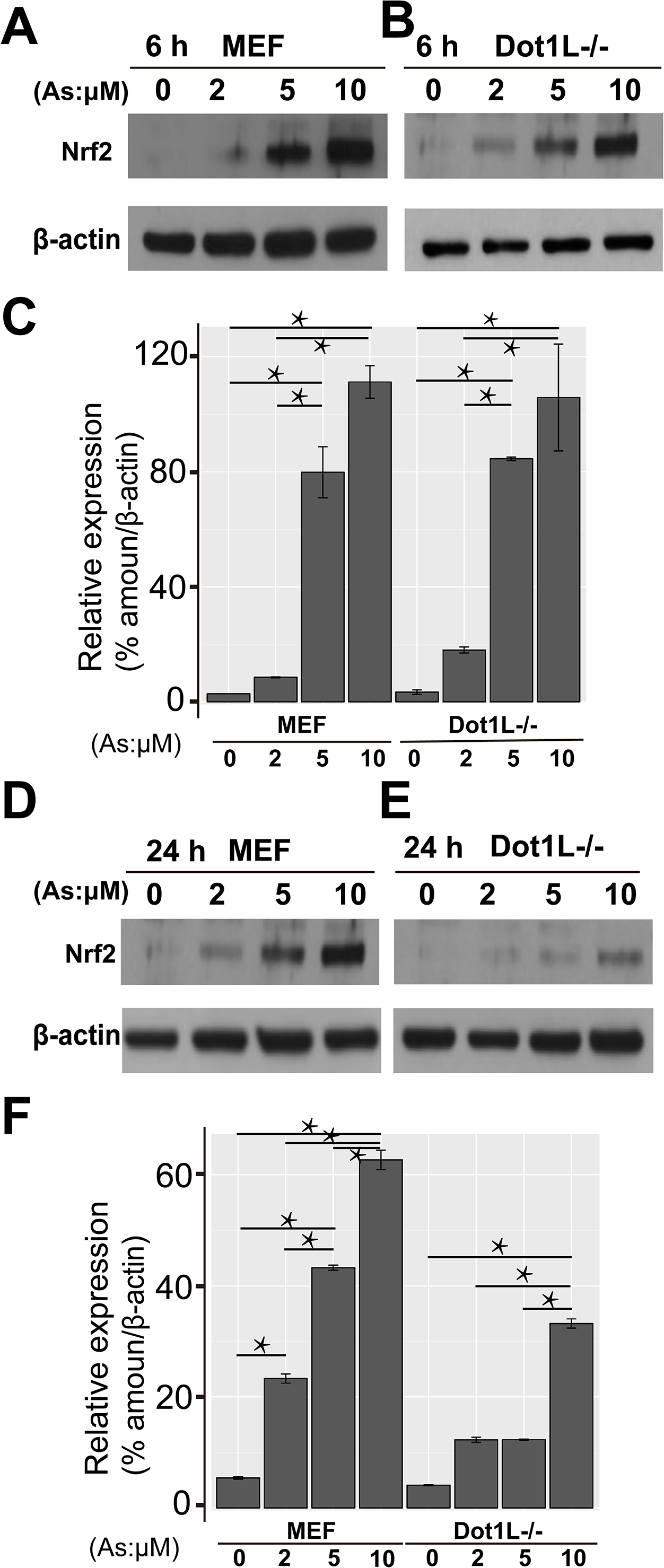
Identification of a low dose exposure condition. 1A and 1B showed results of Western blot of Nrf2 protein in MEF and Dot1L-/-cells after 6 h exposure of arsenite. 1C showed the quantitative results of western blot in 1A and 1B. 1D and 1E showed results of Western blot of Nrf2 protein in MEF and Dot1L-/-cells after 24 h exposure of arsenite. 1F showed the quantitative results of Western blot in. 1D and 1E

### 3.2. Acute low dose exposure of mouse MEF cells for mechanistic insights via RNA-seq analyses

To identify an intermediate factor bridging (external) environmental exposure and (internal) cellular pathway changes, we focused on transcriptomic changes. In other word, we would like to identify the bridging factor(s) based on transcript changes. To our goal, we purified mRNAs from exposed cells and followed our established protocols for RNA-seq library construction (Li *et al.*, 2015). Sequenced reads were aligned to the mouse genome, and expression changes were analyzed. Results demonstrated that arsenite treatment upregulated 518 genes in MEF and 696 genes in Dot1L-/-MEF cells and the overlapping genes were 339 in two cell lines (Fig. 2A). Arsenite treatment also downregulated genes, with 158 genes in MEF and 413 genes in Dot1L-/-MEF cells and the overlapping number was 101 (Fig. 2B). To further validate our RNA-seq results, we selected 10 genes (Up-regulated genes were chosen among genes that played roles in response to oxidative stress and down-regulated genes were chosen among gene that played roles in epigenetics or tumor initiation) for qRT-PCR analyses in both cell lines and confirmed their expression changes (Fig. 2C and 2D). R squares from Pearson correlation analyses were 0.85 for MEF cell and 0.90 for Dot1L-/-cell, suggesting the results from our RNA-seq analyses reliable (Fig. 2E and 2F).

**Fig. 2.**
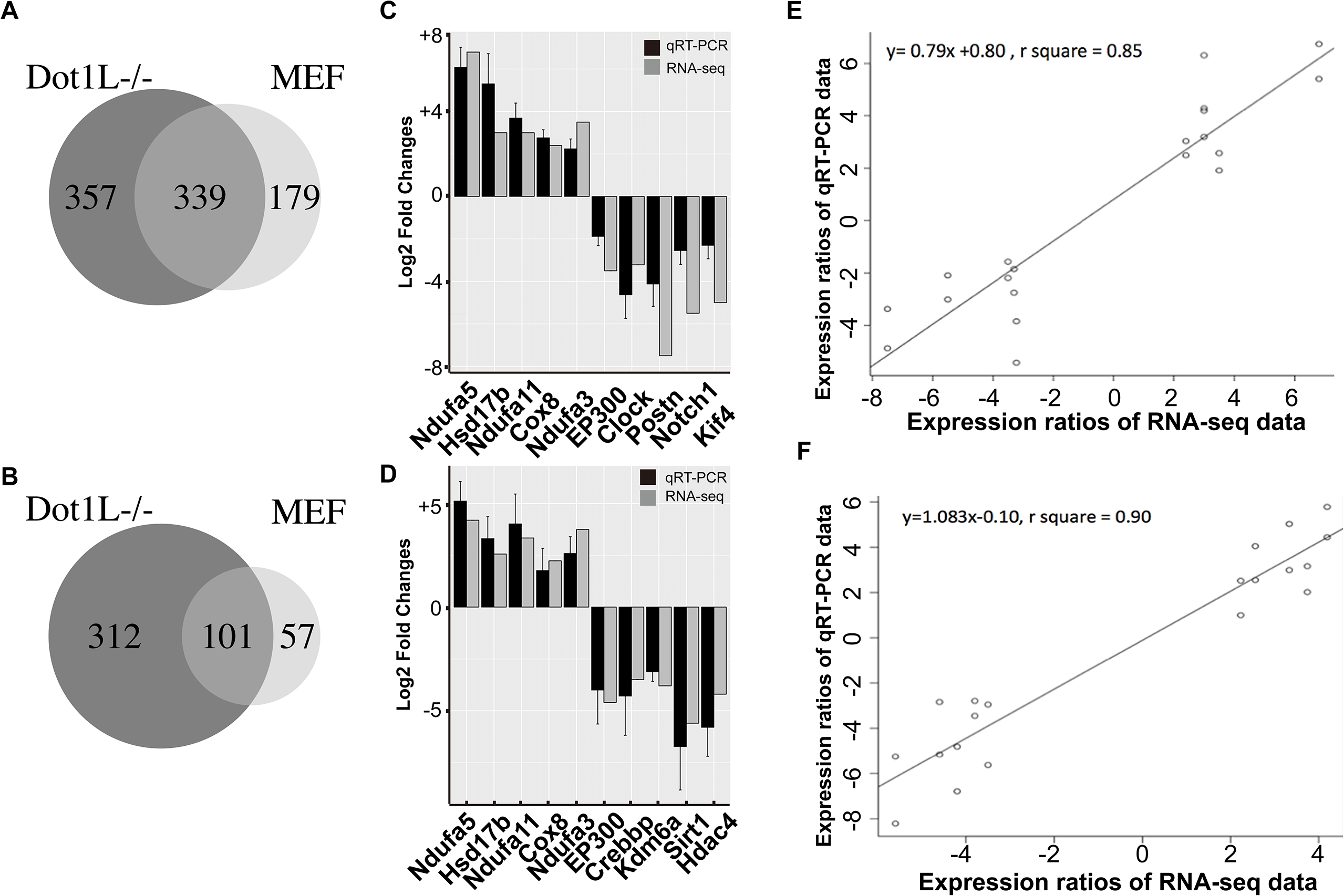
Results from mRNA-seq and qRT-PCR validation. 2A and 2B showed up-regulation and down-regulation genes after arsenite exposure in MEF and Dot1L-/-cells. 2C and 2D showed qRT-PCR results from two cells. 2E and 2F showed Pearson correlation of two cells.

### 3.3. Arsenite exposure inhibited histone acetyltransferase p300

Among these 101 overlapping genes that were inhibited by arsenite treatment in two cell lines, we noticed an intriguing candidate, *EP300* (Log2 fold, -3.5 in MEF cell and -4.6 in Dot1L-/-) (Fig. 2C and 2D). *EP300* encodes histone acetyltransferase p300 (also known as E1A-associated protein p300), essential for regulating cell growth and differentiation and preventing the growth of tumors. For the latter, many have reported the role of p300 as a tumor suppressor (Iyer *et al.*, 2004)

We first did qRT-PCR (Fig. 2C and 2D) and validated the observed changes from RNA-seq analyses. Because reduced RNA level may not reflect the actual protein level and the protein executes enzymatic activity, we next did Western blot to reveal the exact p300 protein level within cells. Western blot from three biological replicates demonstrated that arsenite exposure indeed reduced p300 protein level and showed dose-effect relationships with arsenite concentrations (Fig. 3A and 3B). In combination with RNA-seq and qRT-PCR data, we conclude that arsenite exposure reduces the expression of *EP300* initially at the transcription level and eventually at the protein level.

**Fig. 3.**
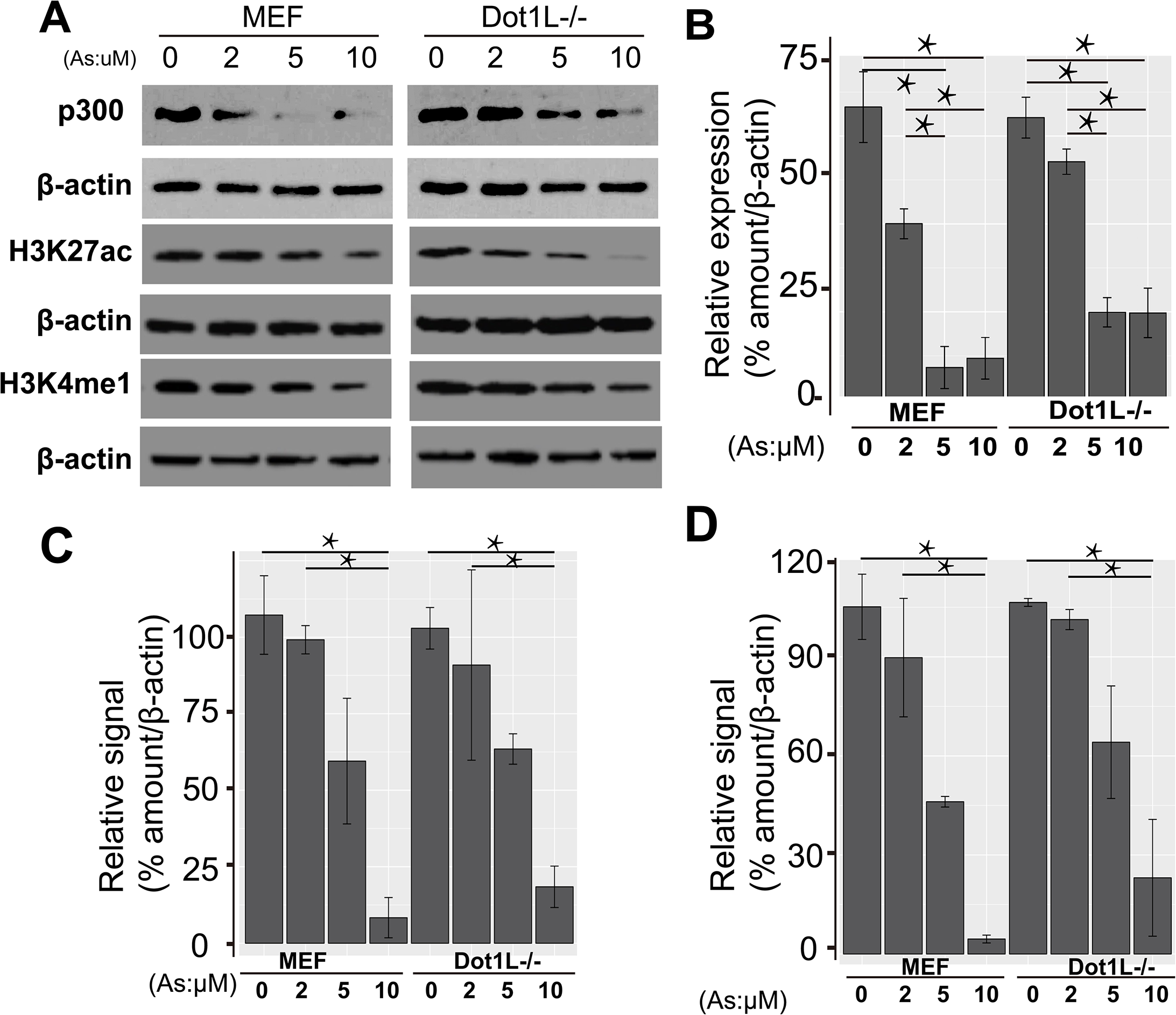
Protein expression and histone modification signals. 3A showed expression of p300, and relative signals of H3K27ac and H3K4me1 in MEF and Dot1L-/-cells in different concentrations of arsenite. 3B, 3C and 3D showed the quantitative results of Western blot.

### 3.4. Altered histone acetylation H3K27ac, an enhancer marker

Next, we determined the consequence of the abolishment of p300 due to arsenite exposure. As a HAT, p300 is expected to acetylate nucleosomal tails. Though p300 seems to acetylate many residues of histone H3 or H4 especially from in vitro assays, there are also reports suggesting H3K27ac as p300-sepcific mark (Li *et al.*, 2011; Tang *et al.*, 2013). Therefore, we examined the changes of H3K27ac after arsenite exposure. With the increasing concentration of arsenite, our data demonstrated that the levels of H3K27ac decreased accordingly (Fig. 3A and 3C) from three replicates. These data demonstrate that arsenite exposure caused decreases of histone H3K27ac, as a consequence of diminished p300.

### 3.5. Arsenite exposure decreased H3K4me1, also a well-known enhancer marker

In epigenetic community, both H3K27ac and H3K4me1 are two well-known markers for enhancers. With H3K27ac down, we asked the extent to which H3K4me1 changes accordingly. To our aim, we checked the signals of H3K4me1 in the same samples. Intriguingly, our Western blot results from three replicates reveal the concomitant decreases of H3K4me1 (Fig. 3A and 3D). These data collectively suggest that arsenite exposure involves underappreciated crosstalk between two enhancer markers, H3K27ac and H3K4me1.

### 3.6. Identification of vulnerable genomic loci with arsenite exposure-reduced H3K27ac

Data of our Western blotting assays reveal the changes of histone acetylation at the global level. To identify the vulnerable genomic loci with altered chromatin after arsenite exposure, we did ChIP-seq analyses using a previously characterized anti-H3K27ac (Wang *et al.*, 2008; Wang *et al.*, 2009). Following our established pipeline, we analyzed H3K27ac ChIP-seq signals genomewide. As expected, arsenite exposed sample has much less H3K27ac peaks compared to control samples. There are 4,772 peaks specific to exposed sample, whereas 18,302 peaks specific to control sample. Two samples shared 21,548 peaks (Fig. 4). These data are consistent with our Western blotting results (Fig. 3).

**Fig. 4.**
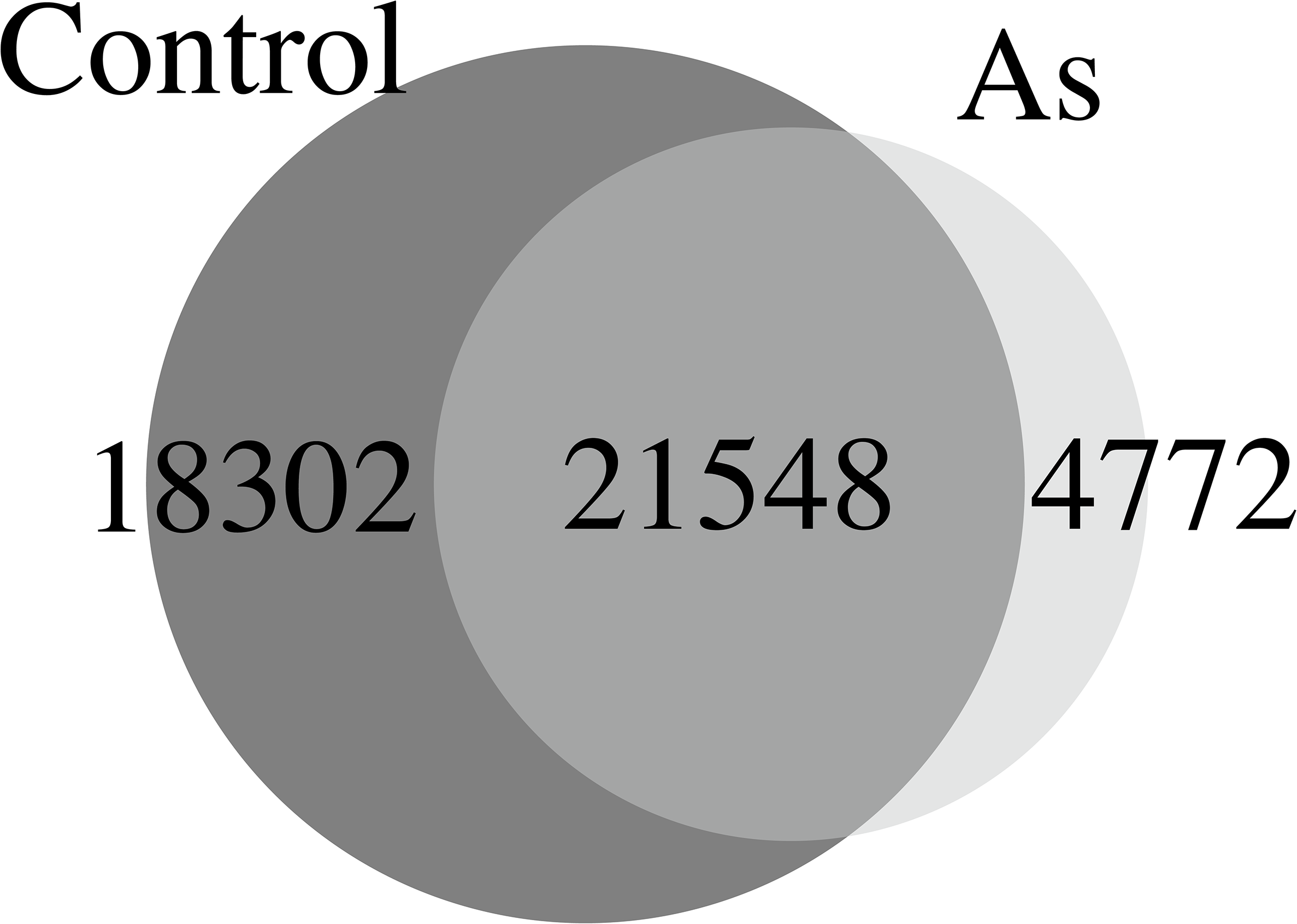
Results from ChIP-seq, which showed peaks in ChIP-seq results using H3K27ac antibody.

### 3.7. Decreased H3K27ac and H3K4me1 inhibited gene expression

Having established that arsenite exposure reduced H3K27ac and H3K4me1 (Fig. 3 and 4), we next determined the reductions on transcription of selected genes. Snapshots of ChIP-seq results were presented for genes including *EP300, Klf4, Notch2*, and *Igf2r* (Fig.5). These data demonstrate that reduction of H3K27ac repressed the transcripts of these genes. Notice the different scales used for presenting RNA-seq data.

**Fig. 5.**
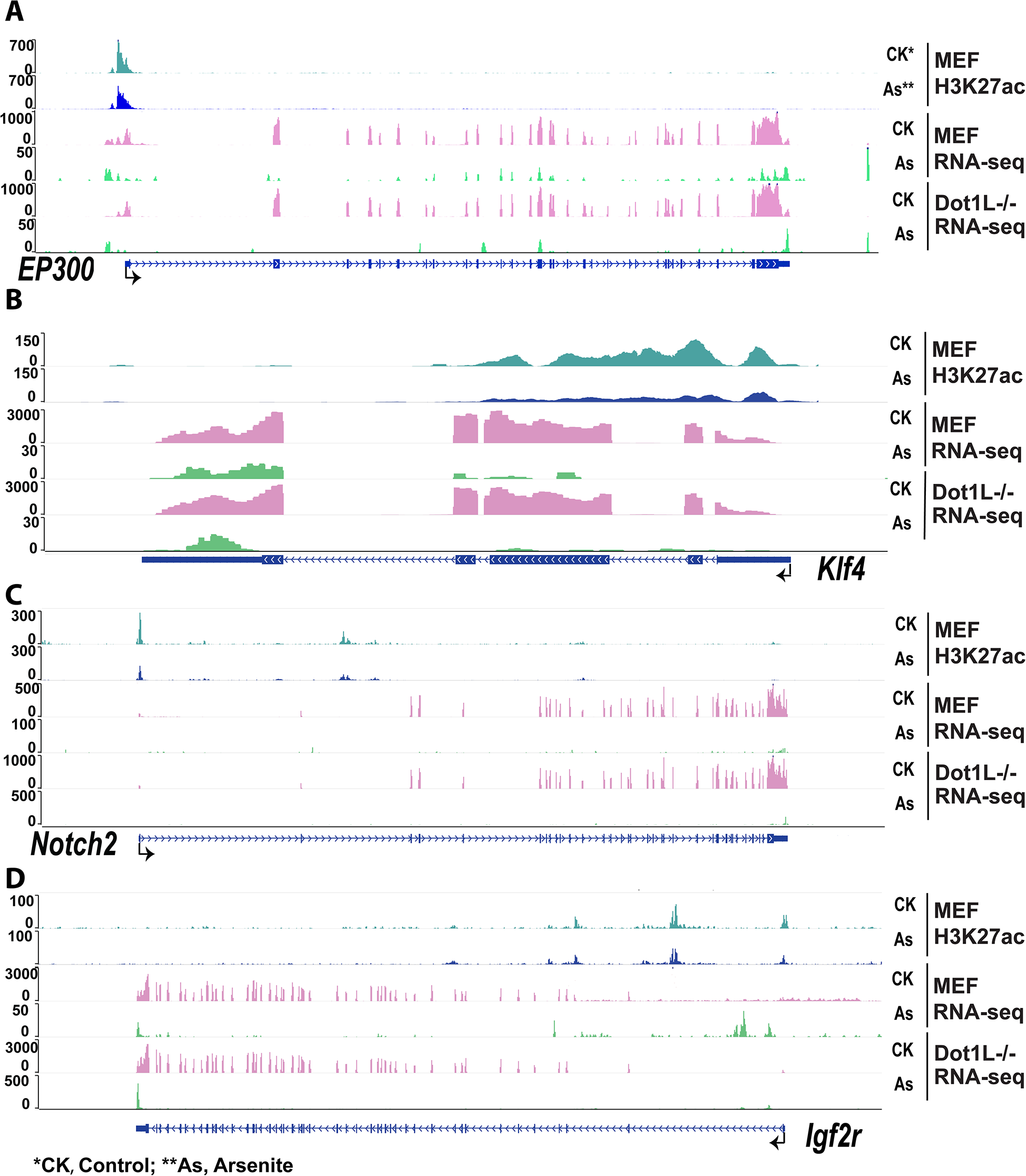
Results from WashU EpiGenome Browser. 6A, 6B, 6C, 6D showed ChIP-seq and mRNA-seq results of *EP300, Klf4, Notch2, and Igf2r* gene.

To provide independent confirmation of H3K27ac at these loci and provide additional information of H3K4me1, we did ChIP-qPCR analyses. Results from Fig. 6 showed the lower enrichment of five gene promoters after arsenic exposure in MEF cell, which were in accordance with ChIP-seq results. *Klf4*, a tumor suppressor gene, showed de-acetylation of H3K27ac and down-regulation (Log2 fold, -5.0), which was validated through ChIP-qPCR (Fig. 3C) after arsenite exposure. Accompany to the decreases of H3K27ac at these loci, our ChIP-qPCR demonstrated the decreases of H3K4me1. These data convinced us that arsenite exposure inhibited p300 protein level within cells, thereby impacting histone markers for enhancers and transcription start sites for gene transcription.

**Fig. 6.**
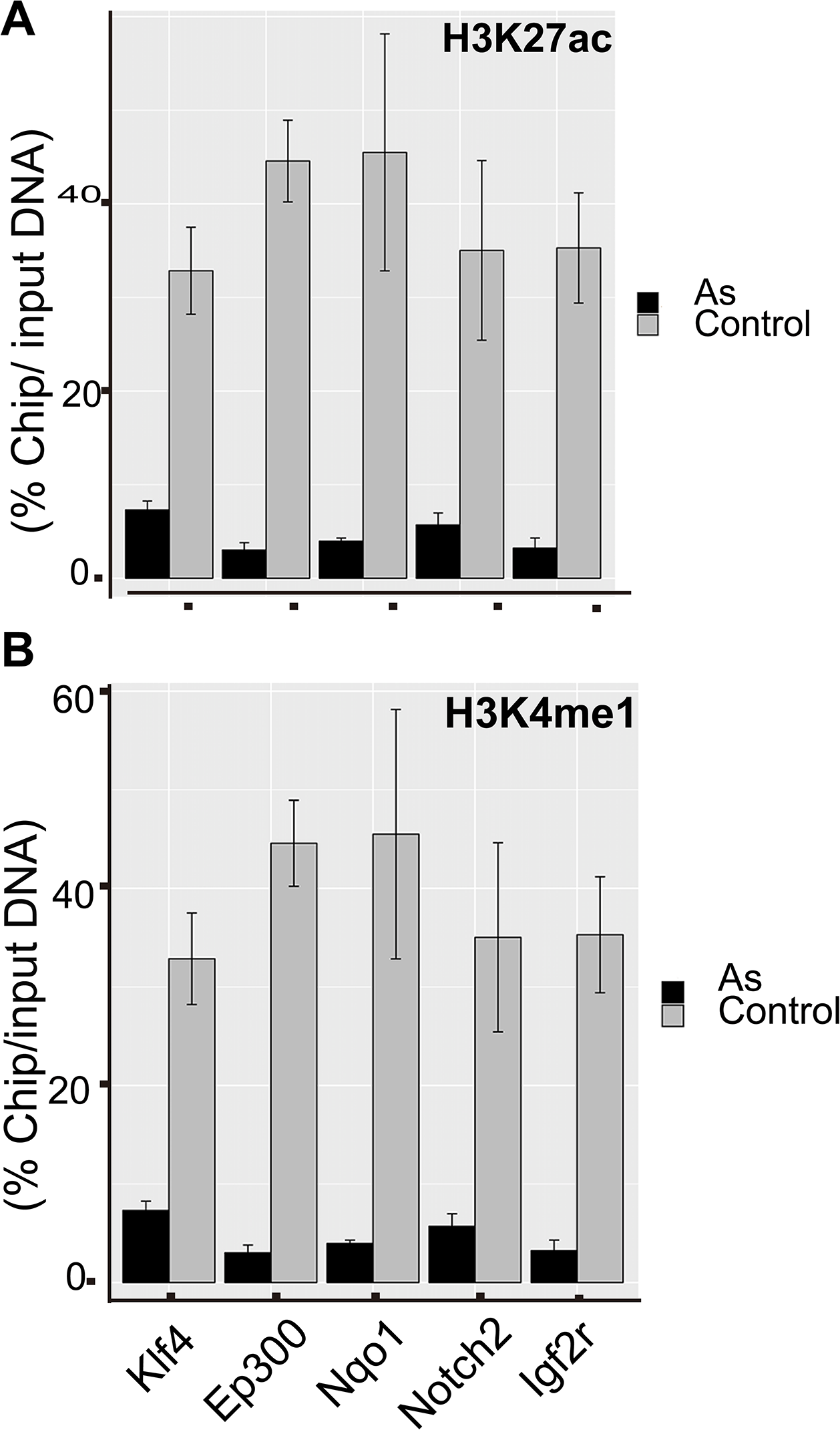
Fig.6 Results from ChIP-qPCR. 7A showed ChIP-qPCR results using H3K27ac antibody and 7B showed ChIP-qPCR results using H3K4me1 antibody.

## 4. Discussion

Considering the expensive nature of deep sequencing and our focus of mechanistic insights to initiate the understanding of epigenomic mechanisms during arsenic exposure, we set out experiments for identification of one low dose condition for future acute exposure. One more reasonable assumption is that exposure at high doses and low doses highly likely shares similar epigenetic mechanism. To our goal, we choose 0, 2, 5, and 10 μM sodium arsenite based on published papers (Shi *et al.*, 2004; Chervona *et al.*, 2012a; Wang *et al.*, 2013; Eckstein *et al.*, 2017) for treatment.

We choose to use MEF cells for a few reasons: First, MEF cell line is a common model used in toxicology. Second, though still very preliminary, H3K79 methylation may be associated with arsenic exposure (Howe *et al.*, 2017). Meanwhile, we have Dot1L knockout MEF cells with depleted histone mark H3K79 dimethylataion (H3K79me2). Therefore, treatments to both MEF and Dot1L-/-help to determine the crosstalk between arsenic exposure and histone H3K79me2 marks in cells. Third, treatments of two independent cell lines would serve as replicates for us. Currently, it is still not conclusive about inorganic arsenic exposure changing histone acetylations for altering gene expression. For example, inorganic arsenic exposure showed H3K27ac and H3K18ac in negative association in females, but positive association in males (Chervona *et al.*, 2012b). Sodium arsenite exposure increased H3K9ac after 24 h exposure in HepG2 cells (Ramirez *et al.*, 2008), which is in dramatic contrast to the decrease of H3K9ac in arsenic exposed mice (Cronican *et al.*, 2013; Pournara *et al.*, 2016). Previous investigations also reported that the chronic arsenic exposure decreased H4K16ac in human UROtsa cells (Jo *et al.*, 2009), via affecting one of the HATs, MYST1 (also known as MOF). MOF forms two distinct multiprotein complexes (MSL and NSL) that both acetylate H4K16ac. Herein, through three biological replicates, we demonstrated that acute arsenic exposure decreased p300 in both mRNA and protein levels (Fig. 2C, 2D, 3A). Consequently, p300-specific H3K27ac was reduced as demonstrated by both ChIP-seq and ChIP-qPCR analyses in our exposed mouse MEF cells. In our system, RNA-seq data did not show the altered expression of MYST1, different from the case in human UROtsa cells. To summarize, three independent investigations including ours showed the decreases of histone acetylations (H3K9ac, H3K27ac, or H4K16ac) (Jo *et al.*, 2009; Cronican *et al.*, 2013), whereas one report showed a contradictory increase of H3K9ac (Ramirez *et al.*, 2008).

Taken together, it remains to be finalized the changes of histone acetylations and affected HAT(s) responsible for the changed histone acetylation upon arsenic exposure. From our observation, all three doses showed similar inhibition of p300 (Figure 3A), maybe ruling out of different doses for different observations (Ramirez *et al.*, 2008). We also noticed the different cell types/lines used or differences in model system used (animal or cell line). Whether these differences explain the controversies of increased or decreased histone acetylation requests further investigation. Lastly, our convincing data suggest that the loss of p300 (and resulting H3K27ac) is related to inorganic arsenic related diseases. Therefore, it may open a new avenue for alleviating the consequence of H3K27ac, for example, via boosting CBP enzymatic activity for at least partially compensating p300 functions (Iyer *et al.*, 2004; Kasper *et al.*, 2006). Both p300 and its regulated gene *Kif4* (silenced after arsenic exposure; Figure 2C) are tumor suppressors(Iyer *et al.*, 2004). For example, *EP300* works as a tumor suppressor in hematological tumors (Oike *et al.*, 1999; Kung *et al.*, 2000). Down-expression of p53 and p300 were also found in azoxymethane treated mice, which is a colon-specific carcinogen(Aizu *et al.*, 2003). The role of *Klf4* as a tumor suppressor has been validated in colorectal cancer (Zhao *et al.*, 2004), non-Hodgkin lymphoma (Guan *et al.*, 2010), classic Hodgkin lymphoma (Guan *et al.*, 2010), and pancreatic ductal carcinoma (Zammarchi *et al.*, 2011).

According to our results, it is reasonable to speculate that arsenite induced down-regulation of p300, and following decreased H3K27ac in *Klf4.* This change led to down expression of *Klf4*, which promoted the initiation of tumors. Other researchers also reported this result (Evans et al., 2007).

## Author Contribution

Z.W conceived this project (with inputs from H.Z, Y.J, and Q.Z) and supervised the experiments. Y.Z, D.L performed the experiments; Y.L did the bioinformatic analyses; D.K, H.Z, Y.J. and Q.Z also conceptually contributed significantly; Y.Z and Z.W wrote the manuscript.

## Acknowledgment

The laboratory of Z.W was supported by U.S. National Institutes of Health (R01ES25761, U01ES026721 opportunity fund, and R21ES028351) and Johns Hopkins Catalyst Award. Q.Z thanks China Scholarship Council for support. This work was also made possible by the ChuTian Professorship from Hubei University, and support from GENEWIZ Suzhou. Lastly, extensive bioinformatic analyses were possible only with computational resources (and/or scientific computing services) at the Maryland Advanced Research Computing Center (MARCC). We are therefore greatly indebted to the support from MARCC.

